# Loss of intestinal endosome associated protein sorting nexin 27 disrupts epithelial barrier and promotes inflammation

**DOI:** 10.1101/2025.02.05.635995

**Authors:** Shreya Deb, Yongguo Zhang, Yinglin Xia, Jun Sun

## Abstract

**Backgrounds and aims:** SNX27, member of the sorting nexin (SNX) family, carries a unique PDZ domain and mediates recycling of endocytosed transmembrane proteins. SNX27 is critical for neurodevelopmental processes, however its role in intestine remains unexplored. We aim to determine the previously unknown roles of SNX27 in regulating intestinal homeostasis, epithelial barrier integrity, and inflammatory responses.

**Methods:** We used available datasets to analyze SNX27 expression in human IBD. We generated a novel mouse model of SNX27 conditional deletion from intestinal epithelial cells (SNX27^ΔIEC^) and challenged these mice with Dextran Sulfate Sodium (DSS).

**Results:** SNX27 expression was significantly lower in human IBD, including UC and CD. SNX27^ΔIEC^ mice had significantly lower bodyweight and exhibited increased proliferation and poor differentiation of secretory Paneth and Goblet cells. We found reduced mucin layer and downregulation of crucial epithelial barrier proteins Δ-catenin, E-cadherin, ZO-1, and Claudin10 in SNX27^ΔIEC^ mice. SNX27^ΔIEC^ mice showed high intestinal permeability and spontaneously developed intestinal inflammation. Moreover, SNX27^ΔIEC^ mice were more susceptible towards DSS-induced colitis, compared to the SNX27^Loxp^ mice.

**Conclusion:** Overall, deletion of intestinal epithelial SNX27 weakens barrier functions and promotes inflammation. Our results indicate a novel role of SNX27 in regulating intestinal physiology and protecting against intestinal disorders. Thus, understanding the mechanisms of SNX27 downregulation in IBD will provide insights into new prevention and targets against chronic inflammation.

**Synopsis:**

- SNX27 recycles internalized transmembrane proteins in the endocytic pathway. Human IBD showed reduced levels of SNX27.
- SNX27 plays novel functions by maintaining intestinal homeostasis and inhibiting inflammation.
- SNX27 protects the host against losing intestinal integrity during inflammation.

## Introduction

Sorting nexin (SNX) proteins play fundamental roles in regulating intracellular trafficking and sorting of internalized cargo in the endocytic pathway ^1^. SNX27 is a unique member of this family of 33 proteins because it carries a third PDZ (post-synaptic density 95/discs large/zonula occludens-1) binding domain, in addition to the PX (conserved) and FERM domains^2,3^. This makes SNX27 a distinct sorting protein at the structural as well as functional level. SNX27 is known to mediate the rescue of endocytosed transmembrane proteins from lysosomal degradation in a PDZ-domain dependent manner ^3^. Therefore, defects in the SNX27-dependent endosomal recycling pathway leading to the improper sorting of internalized cargo have been linked to a variety of human diseases ^4–6^. The SNX27 was first identified in the rat neocortex by Joubert et al. in 2004 and since then SNX27 has been reported to be associated with several critical neurological diseases, e.g., Down’s Syndrome (DS) and Alzheimer’s Disease (AD) ^5,7,8^,. SNX27-mediated recycling of cell surface protein β2-adrenergic receptor (β2AR) has been implicated in the regulation of immune cell function and inflammatory signaling pathways ^3^. Moreover, SNX27 has been shown to interact with and modulate the activity of several key signaling proteins involved in AKT and ERK pathways, which play critical roles in regulating inflammatory responses ^9^ and tumorigenesis ^10^. However, SNX27 in the maintenance of the intestinal homeostasis remains largely unexplored.

Inflammatory bowel disease (IBD) ^11^ is an autoimmune disorder that affects the gastrointestinal tract and can be categorized into Crohn’s disease ^12^ and ulcerative colitis (UC), depending on the site of inflammation ^13^. The etiology of IBD has been characterized as chronic inflammation resulting from a combination of several factors, including host genetics, environmental factors, micronutrients, and microbiome ^11,14,15^. Integrity of the intestinal epithelial barrier plays an essential role in maintaining tissue health by preventing unwanted invasion of pathogens and toxins contributing to mucosal inflammation ^16^. This is strictly regulated by a complex of barrier proteins that form the tight/adherens junction (TJ/AJ) and dysregulation in their expression, and localization may contribute to IBD ^16^. For example, an increase in the TJ protein Claudin2 and a decrease in Claudin5 were shown to promote leaky gut barrier in IBD patients ^17^. Many of these TJ/AJ proteins carry a PDZ-binding motif and undergo endogenous endocytosis and recycling which implicates towards their interactions with SNX27 ^18,19^. However, the role of SNX27 in regulating intestinal epithelial barrier integrity has not been investigated *in vivo*, while the association of endosomal sorting and recycling in intestinal inflammation is still poorly understood.

In the current study, we focus on addressing two fundamental questions: (1) is SNX27 involved in human IBD, and (2) how does SNX27 contribute to chronic inflammation? We found that SNX27 expression is downregulated in UC and CD patients. Previous total body knockouts of SNX27 in mice have been embryonically lethal ^20^. Therefore, we generated a novel conditional knockout mouse model of SNX27 harboring a tissue-specific deletion from the intestinal epithelial cells (IECs) only for *in vivo* studies. We then investigated the impact of intestinal epithelial SNX27 on barrier integrity by regulating epithelial junctions. We also examine SNX27 regulation of Paneth cell and Goblet cell function, which are essential for protecting the intestinal epithelium from luminal insults and maintaining mucosal homeostasis. By investigating the mechanisms underlying the impact of intestinal epithelial SNX27 deletion, we aimed to elucidate the previously unknown roles of SNX27 in the pathogenesis of IBD.

## Results

### Downregulated expression of SNX27 in patients with IBD

To determine the clinical relevance of SNX27 in human IBD, we analyzed publicly available datasets in the NCBI GEO database (https://www.ncbi.nlm.nih.gov/geo). We studied the expression levels of SNX27 in patients with either UC or CD and compared them to healthy controls. The dataset with accession number GSE6731 belongs to a study which used high-density oligonucleotide microarrays to examine the gene expression levels between 13 UC, 19 CD and 4 healthy patients ^21^. Upon reanalyzing this dataset, we found that the gene expression levels of SNX27 in the colons were significantly downregulated in patients with either UC or CD, compared with non-IBD patients **(Fig. 1A)**. Similarly, we analyzed another dataset with accession number GSE11223, which analyzed colonic epithelial biopsies of various anatomic locations from 67 patients with UC and 31 healthy controls ^22^. This study harvested tissues that were histologically deemed unaffected vs actively inflamed. We also observed a significant downregulation of SNX27 gene expression in patients with UC vs healthy controls, with a significantly lower expression exhibited in tissues with active inflammation compared to uninflamed tissues within the same cohort of patients with UC **(Fig. 1B)**. Further analysis of GSE11223 revealed varying expression levels of SNX27 in different regions of the colon **(Fig. 1C)**. We observed that SNX27 is highly expressed in the descending colon and least in the sigmoid colon of both healthy and UC affected patients, with progressive downregulation observed in inflamed tissues, as compared to normal tissues **(Fig. 1C)**. Overall, our analysis indicates that SNX27 is significantly downregulated upon inflammation and induction of human IBD.

**Figure 1.**
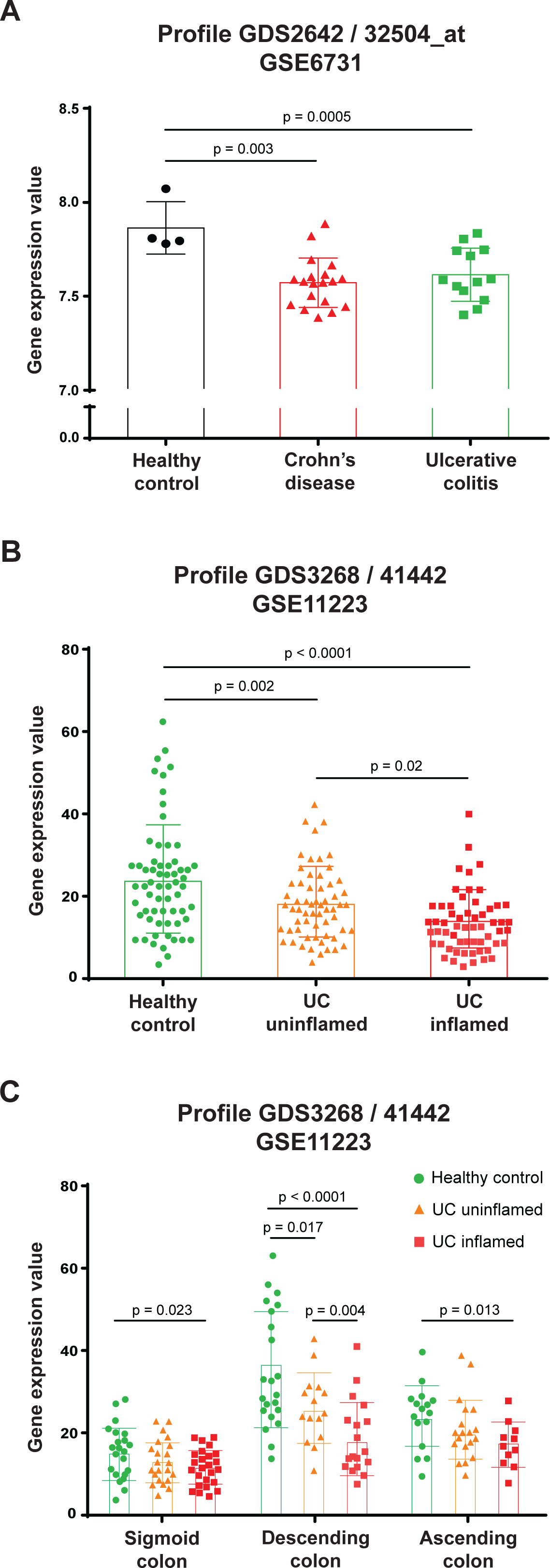
Reduced SNX27 expression in human IBD. **(A)** SNX27 expression was significantly downregulated in CD or UC patients, compared with non-IBD patients. The data were retrieved from National Center for Biotechnology Information (NCBI) GEO database with accession number GSE6731. Data are shown as mean ± SD, one-way ANOVA test, healthy controls, n = 4; CD patients, n = 19; UC patients, n = 13. **(B)** GSE11223 database also revealed significantly lower SNX27 expression in patients with UC (uninflamed and inflamed section) vs healthy controls. Furthermore, SNX27 expression was significantly lower in tissues with active inflammation compared to uninflamed tissues within the same cohort of UC patients. Each dot on the graph represents a single tissue sample; multiple samples from various regions of the colon were collected from individual subjects. Data are shown as mean ± SD, one-way ANOVA test, healthy controls, n = 31; UC patients, n = 67. **(C)** SNX27 is highly expressed in the descending colon and least in the sigmoid colon of both healthy and UC affected patients, with progressive downregulation observed in inflamed tissues, as compared to normal tissues in GSE11223. Each dot on the graph represents a single tissue sample; multiple samples from various regions of the colon were collected from individual subjects. Data are shown as mean ± SD, two-way ANOVA test, healthy controls, n = 31; UC patients, n = 67. All *p* values are shown in the figures.

### Establishing a novel mouse model with a conditional deletion of SNX27

To study the fundamental role of SNX27 in the intestines, we designed a conditional knockout mouse model of SNX27 harboring a tissue-specific deletion from the intestinal epithelial cells (IECs). To our knowledge, our lab is the first one to report this IEC-specific SNX27 knockout in mice. We used transgenic mice expressing cre recombinase driven under the promoter region of mouse Villin gene, only expressed in IECs of the small and large intestine, to cross with the SNX27^Loxp^ mice **(Fig. 2A)**. We checked for the mRNA expression levels of SNX27 in the mouse colon and ileum tissues and observed a significant downregulation in SNX27^ΔIEC^ mice **(Fig. 2B)**. We further validated the deletion at the protein level via western blot using intestinal mucosal scraping and observed a significant downregulation of SNX27 protein levels in colon as well as ileum of SNX27^ΔIEC^, compared to SNX27^Loxp^ mice **(Fig. 2C)**. We examined the localization of SNX27 and visualized the conditional knockout by performing immunohistochemical staining of mice proximal colons. Our analysis revealed a significant downregulation of SNX27 protein expression levels in SNX27^ΔIEC^ mice which was visually confirmed by the lack of SNX27-specific DAB staining at the top of the colonic crypts, whereas the SNX27^Loxp^ mice express SNX27 at the base as well as the top of the colonic crypts **(Fig. 2D)**. Therefore, we confirm successful deletion of SNX27 from IECs in the SNX27^ΔIEC^ mouse model.

**Figure 2.**
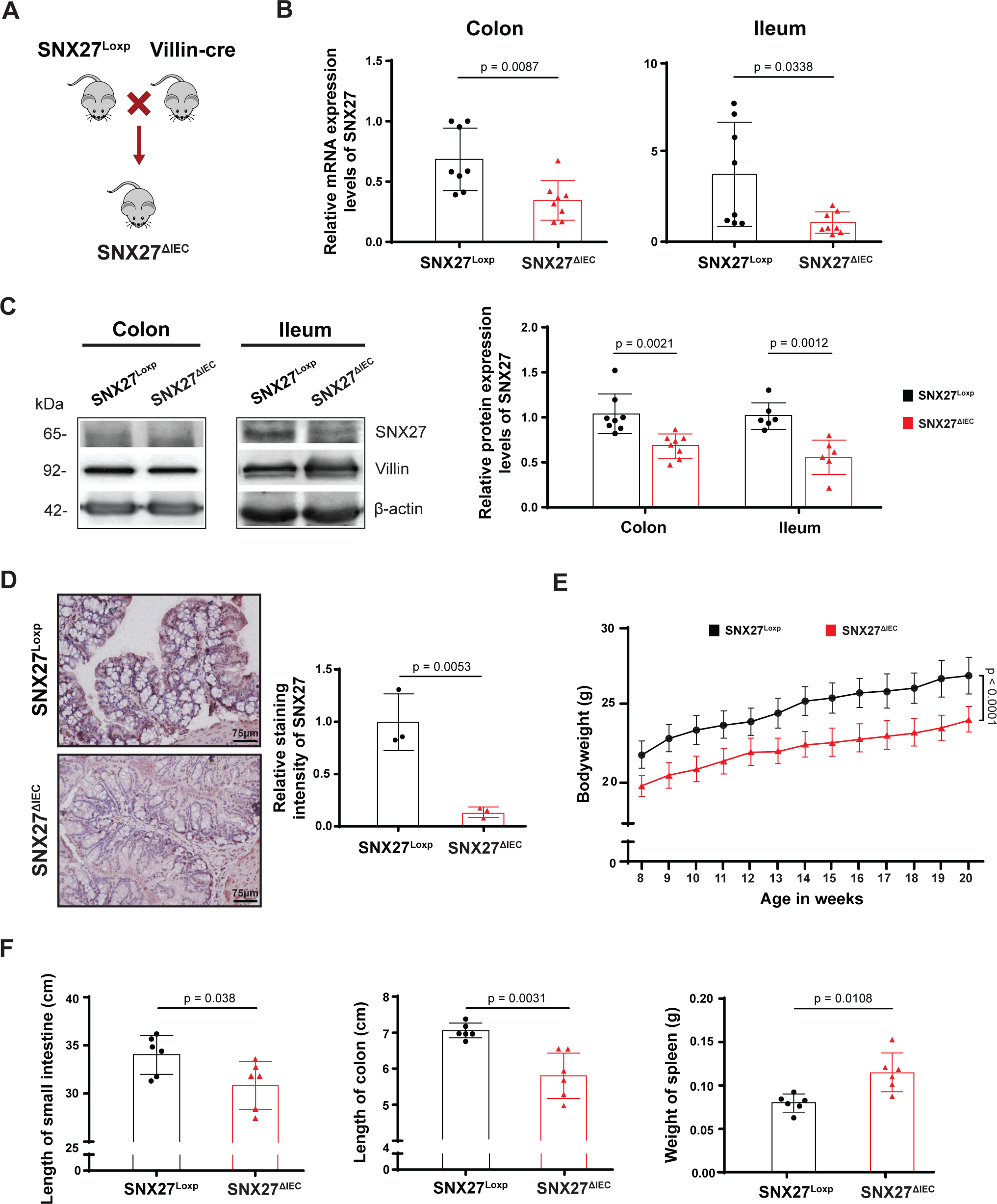
Establishment of SNX27^ΔIEC^ mouse model. **(A)** SNX27^ΔIEC^ mice were developed by crossing the SNX27^Loxp^ mice with IECs-specific Villin-cre expressing mice that resulted in a tissue-specific conditional knockout of SNX27 from small and large intestine. **(B)** mRNA and **(C)** protein expression levels of SNX27 in mice colon and ileum tissues were significantly downregulated in SNX27^ΔIEC^ mice compared to SNX27^Loxp^ mice. All data are expressed as mean ± SD, Welch’s *t*-test, n = 8. **(D)** Immunohistochemistry (IHC) staining of colon tissues further validated loss of SNX27 from IECs in SNX27^ΔIEC^ mice. The relative IHC intensity was quantified with ImageJ. Data are shown as mean ± SD, Welch’s *t*-test, n = 3. **(E)** SNX27^ΔIEC^ mice had a significantly lower body weight compared to SNX27^Loxp^ mice during the various stages of development ranging from 8 to 20 weeks old. Data are shown as mean ± SEM, two-way ANOVA test, n = 11. **(F)** SNX27^ΔIEC^ mice had significantly shorter small intestine, colon, and larger spleen, compared to SNX27^Loxp^ mice. All data are expressed as mean ± SD, Welch’s *t*-test, n = 6. All *p*-values are shown in the figures.

### SNX27 expression regulates intestinal tissue physiology in mice

With our newly generated SNX27^ΔIEC^ mouse model, we were able to examine the physiological changes at the basal level upon loss of intestinal SNX27. We observed that the SNX27^ΔIEC^ mice had a significantly lower bodyweight compared to SNX27^Loxp^ mice during the various stages of development (from 8 to 20 weeks old) **(Fig. 2E)**. We measured intestinal length and observed that SNX27^ΔIEC^ mice had significantly shorter small intestines and colons compared to SNX27^Loxp^ mice (**Fig. 2F)**. These body weight changes signify underlying abnormalities and/or illness, as improper absorption of nutrients is known to impact growth and development ^23^. Shortening of the gastrointestinal tract often results in malabsorption of food and fluids due to the lack of available surface area and is a known characteristic of IBD ^24^.

Additionally, we compared weight of spleen and found that SNX27^Loxp^ mice exhibited significantly enlarged spleens **(Fig. 2F)**.

### SNX27 deletion alters intestinal epithelial cell (IEC) proliferation and apoptosis

Intestinal homeostasis is highly dependent on the proper functioning and distribution of IECs, the major component shaping the gut physiology ^25^. Therefore, we analyzed the effects of SNX27 deletion on the proliferative capacity of IECs by using Proliferating Cell Nuclear Antigen (PCNA) immunofluorescence staining. There was a significant upregulation of PCNA positive cells along the base of the crypts in both colon and ileal tissues of SNX27^ΔIEC^ mice **(Fig. 3A)**. As the balance between epithelial cell proliferation and apoptosis in the intestinal mucosa is a tightly regulated process, any changes in this dynamic can disrupt the fate of intestinal homeostasis ^26^. We next looked into apoptosis, a critical component that determines IEC turnover ^26^. We observed a significant upregulation in the number of TUNEL-positive apoptotic cells in SNX27^ΔIEC^ mice colons, compared to the SNX27^Loxp^ mice **(Fig. 3B)**. Most apoptotic cells were detected in the apical side in the colon of SNX27^Loxp^ mice, whereas apoptotic cells were identified from the apical to the basolateral side of colon in SNX27^ΔIEC^ mice. We further validated the induction of apoptosis by performing western blot analysis of expression levels of the pro-apoptotic protein cleaved or active caspase-3 and anti-apoptotic protein Bcl-xL. There was a significant upregulation of active caspase-3, while Bcl-xL was significantly downregulated, in the colonic epithelial cells of SNX27^ΔIEC^ vs SNX27^Loxp^ mice **(Fig. 3C)**.

**Figure 3.**
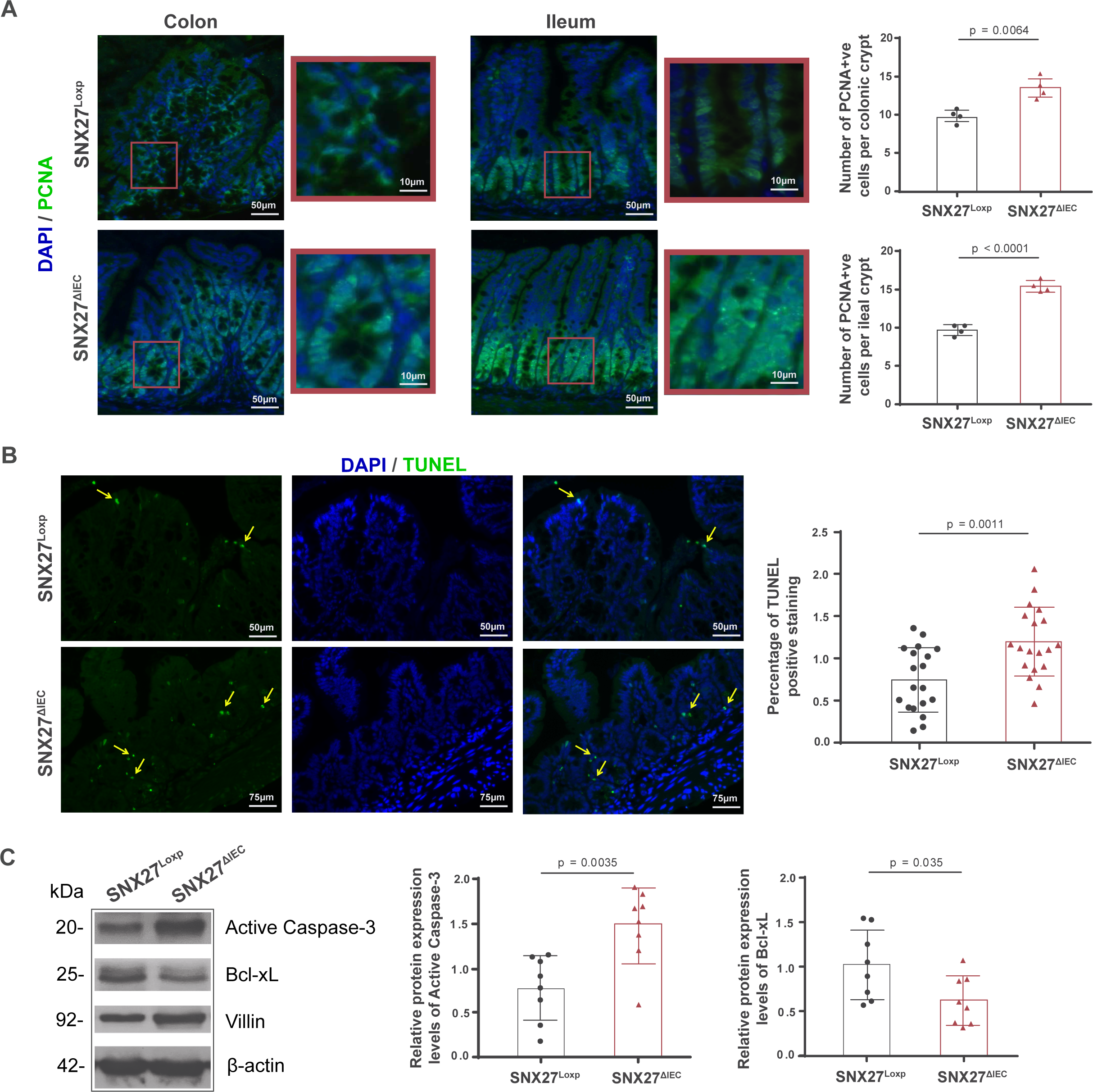
Loss of intestinal SNX27 promoted crypt cell proliferation and IEC apoptosis. **(A)** Immunofluorescence (IF) staining of PCNA in colon and ileum tissues showed greater number of PCNA positive cells per crypt in SNX27^ΔIEC^ mice. All data are expressed as mean ± SD, Welch’s *t*-test, n = 4. **(B)** TUNEL staining of colon tissues showed significant upregulation in the number of apoptotic cells in SNX27^ΔIEC^ mice. All data are expressed as mean ± SD, Welch’s *t*-test, n = 19. **(C)** SNX27^ΔIEC^ mice had increased active capase-3 and lower Bcl-xL protein levels compared to SNX27^Loxp^ mice. All data are expressed as mean ± SD, Welch’s *t*-test, n = 8. All *p*-values are shown in the figures.

### SNX27 deletion altered numbers of Paneth cells and Goblet cells

Some IECs differentiate into secretory lineage, particularly Paneth cells (PC) and Goblet cells (GC). PCs are predominantly found in the small intestine and play a critical role in regulating host immune responses and shaping the commensal gut microbiome ^27^. We analyzed PC count and phenotype based on the localization of lysozyme staining, using a previously described method ^28^. We found an overall downregulation in the number of PCs in the ileal crypts of SNX27^ΔIEC^ mice. Among these, we observed a significant upregulation in the number of abnormal PCs, as compared to SNX27^Loxp^ mice **(Fig. 4A)**.

**Figure 4.**
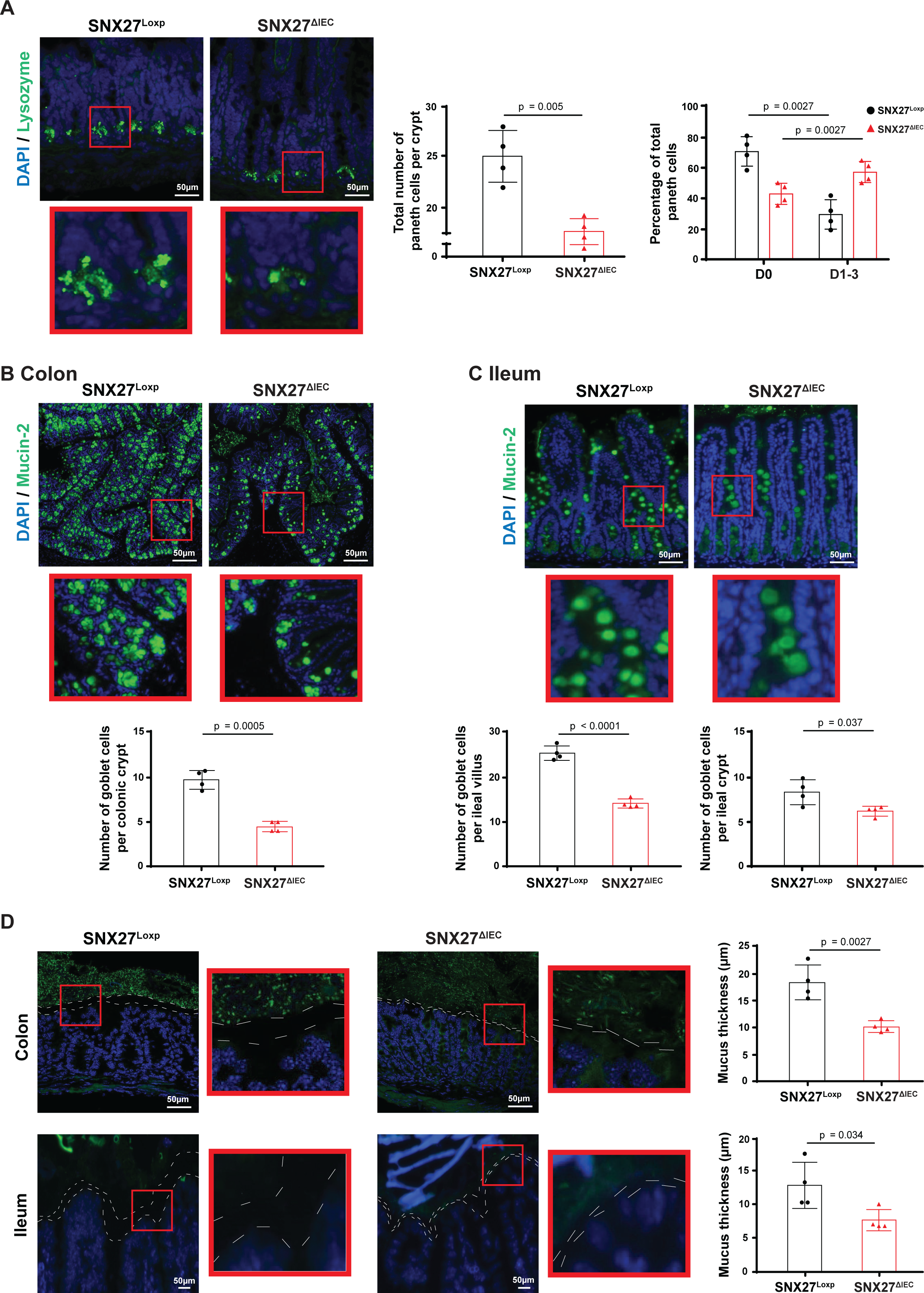
Deletion of SNX27 promoted loss of Paneth and Goblet cells. **(A)** SNX27^ΔIEC^ mice had a significantly lower number of PCs in the ileal crypts, with an increase in abnormal phenotype. Normal PCs were indicated as D0, while abnormal PCs were grouped as D1 (disordered), D2 (depleted), and D3 (diffused) lysozyme granule morphologies. All data are expressed as mean ± SD, Welch’s *t*-test or two-ANOVA test respectively, n = 4. **(B)** and **(C)** Mucin-2 staining used to determine goblet cell distribution in colon and ileum showed significant downregulation in SNX27^ΔIEC^ mice compared to SNX27^Loxp^ mice. All data are expressed as mean ± SD, Welch’s *t*-test, n = 4. **(D)** FISH staining was used to visualize the mucus thickness and showed significant narrowing of the mucus layer in SNX27^ΔIEC^ mice compared to SNX27^Loxp^ mice. All data are expressed as mean ± SD, Welch’s *t*-test, n = 4. All *p*-values are shown in the figures.

GCs are another crucial component of the secretory lineage that maintain intestinal homeostasis by producing mucus that separates the host tissue from luminal gut contents by creating a protective mucin layer ^29^. There was a significant downregulation in Muc2 positive GCs in both colonic and ileal tissues of SNX27^ΔIEC^ vs SNX27^Loxp^ mice **(Fig. 4B and Fig. 4C)**. Proper number and distribution of GCs is important to maintain the mucin layer which acts as an intestinal barrier, in addition to epithelial cells and microbiome, to prevent harmful toxins and pathogens from entering the mucosa and causing inflammation ^29^ ^26^. We performed Fluorescence in situ hybridization (FISH) ^30^ using antisense single-stranded DNA probes targeting the bacterial 16S ribosomal RNA. This probe, EUB338 (5′-GCTGCCTCCCGTAGGAGT-3′), conjugated to Alexa Fluor488 probe allows to visualize the microbiome as the 16S ribosomal RNA is highly conserved between different species of bacteria. We observed reduced bacteria abundance (green staining) in the SNX27^ΔIEC^ mice, compared to SNX27^Loxp^ mice **(Fig. 4D)**. This method also allows us to measure the mucus thickness in the intestinal tissue and we observed a significant narrowing of the mucus layer in mice lacking the intestinal SNX27 **(Fig. 4D)**.

### Lack of intestinal SNX27 damages epithelial barrier by altering tight junction proteins

Tight-junction (TJ) proteins play a critical role in holding the individual IECs together to maintain the architecture and integrity of the intestinal epithelial barrier ^18^. TJ protein Zonula Ocludens-1 (ZO-1) directly interacts with other TJ proteins via its PDZ domain and binds them to the actin cytoskeleton, thereby maintaining their expression at the epithelial junctions ^19^. Through immunofluorescence staining, we observed that ZO-1 expression levels were significantly diminished at the epithelial barrier in SNX27^ΔIEC^ mice **(Fig. 5A)**. This indicates dysregulation in the arrangement of TJ proteins with loss of intestinal SNX27. To further validate the weakening of the epithelial barrier, we analyzed expression levels of Claudin10, a member of the Claudin family of TJ proteins. Immunofluorescence staining in mice colon tissues showed significant downregulation of Claudin10 at the apical membrane of SNX27^ΔIEC^ mice **(Fig. 5B)**. We also validated these observations via western blot analysis in the scraped colon epithelium of SNX27^ΔIEC^ vs SNX27^Loxp^ mice which showed similar results **(Fig. 5C)**. Therefore, SNX27^ΔIEC^ mice exhibit reduction of the TJ proteins, e.g., ZO-1 and Claudin10.

**Figure 5.**
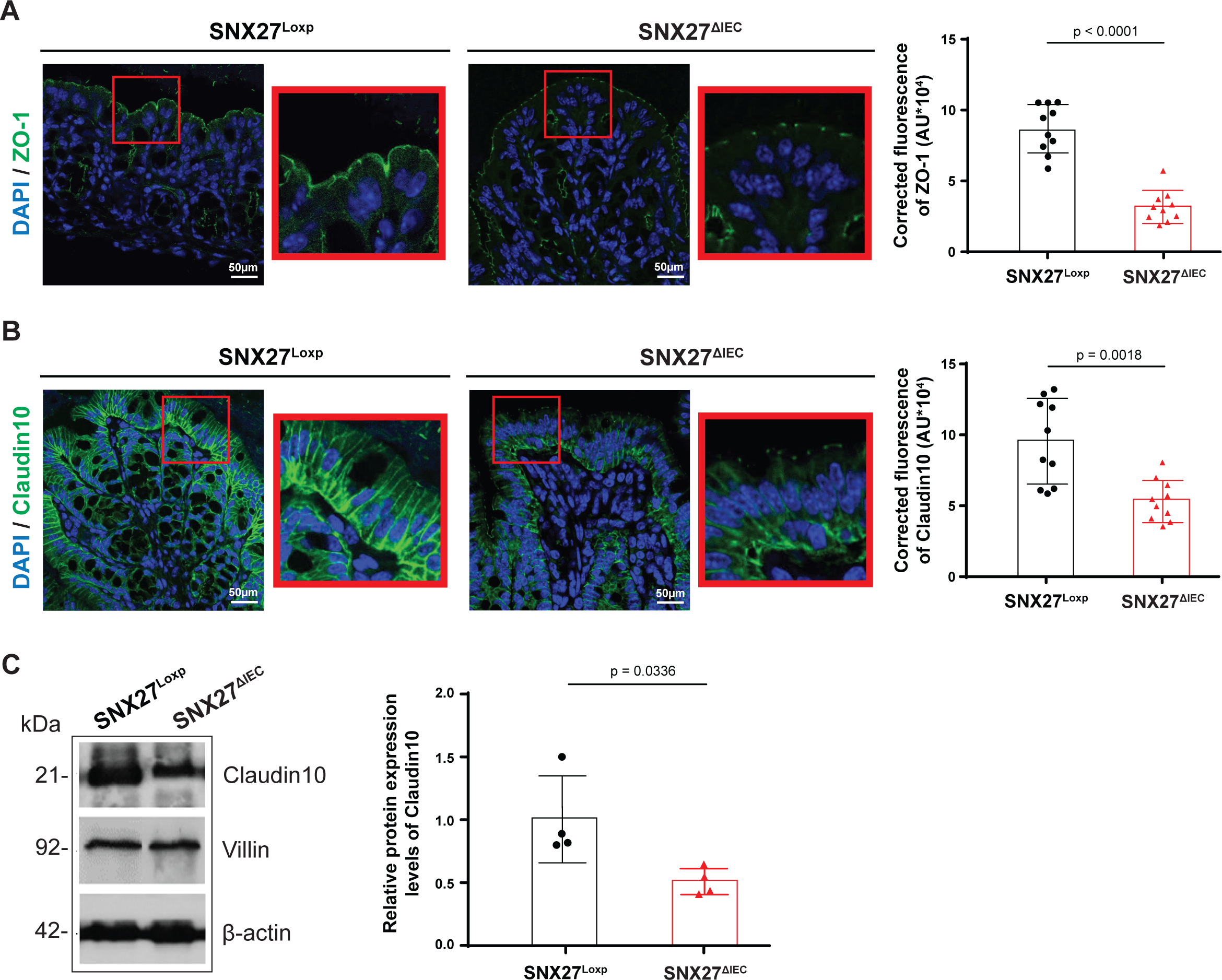
Lack of intestinal SNX27 disrupted epithelial barrier by altering expression levels and localization of TJ proteins. **(A)** ZO-1 expression levels were disrupted and decreased at the epithelial barrier in SNX27^ΔIEC^ mice compared to SNX27^Loxp^ mice by IF staining. Data are expressed as mean ± SD, Welch’s *t*-test, n = 10. **(B)** SNX27^ΔIEC^ mice had significantly downregulation Claudin10 expression compared to SNX27^Loxp^ mice by IF staining. Data are expressed as mean ± SD, Welch’s *t*-test, n = 10. **(C)** Protein expression levels of Claudin10 in colonic epithelial cells was downregulated in SNX27^ΔIEC^ mice compared to SNX27^Loxp^ mice. Data are expressed as mean ± SD, Welch’s *t*-test, n = 4. All *p*-values are shown in the figures.

### SNX27 deletion promotes gut leakiness and inflammatory responses

An increase in gut leakiness has been associated with disruption of intestinal barriers and susceptibility to IBD ^26^. Therefore, we compared intestinal permeability levels between SNX27^ΔIEC^ and SNX27^Loxp^ mice by analyzing serum levels of FITC-Dextran, 4 hours after oral gavage. SNX27^ΔIEC^ mice had significantly higher concentrations of FITC-Dextran in the serum, indicating that intestinal deletion of SNX27 contributes towards increased leakiness of the gut **(Fig. 6A)**. Consequently, serum concentration levels of lipopolysaccharides (LPS), a gram-negative bacteria endotoxin, was found to be significantly higher in SNX27^ΔIEC^ mice **(Fig. 6B)**. High intestinal permeability levels and serum LPS detection both drive systemic mucosal inflammation ^26^. Histological analysis of SNX27^ΔIEC^ colons revealed a significantly higher inflammation score, determined by an observable increase in inflammatory cell infiltration and crypt architecture distortion **(Fig. 6C)**. Therefore, we analyzed tissue cytokine levels via qPCR analysis and observed that mRNA levels of proinflammatory markers TNF-α and IL-6 were significantly higher in the colons of SNX27^ΔIEC^ mice, compared to SNX27^Loxp^ mice **(Fig. 6D)**.

**Figure 6.**
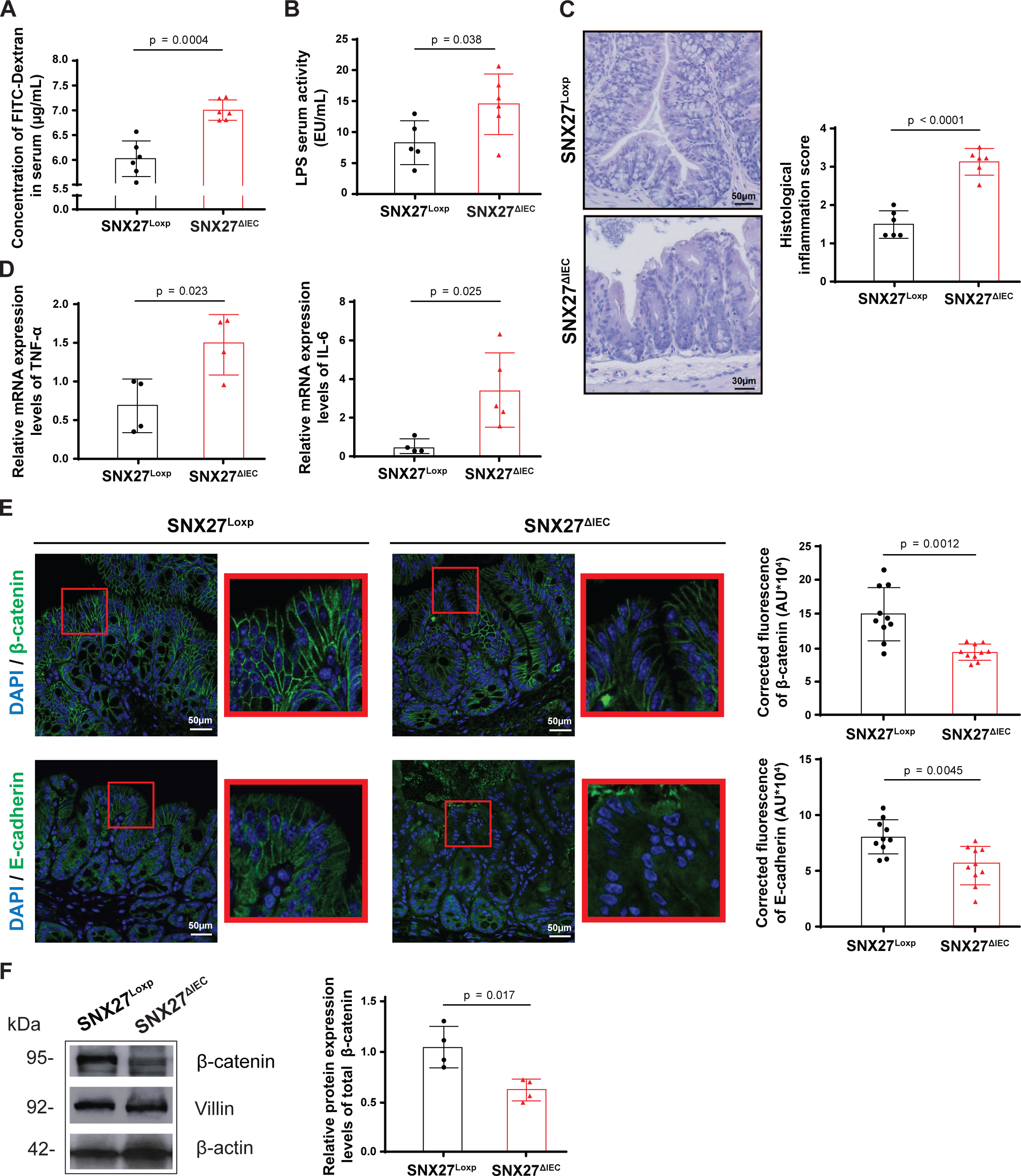
SNX27^ΔIEC^ mice had increased intestinal permeability and chronic inflammation. **(A)** SNX27^ΔIEC^ mice had significantly higher intestinal permeability levels compared to SNX27^Loxp^ mice. All data are expressed as mean ± SD, Welch’s *t*-test, n = 6. **(B)** Loss of intestinal SNX27 also promoted serum LPS concentration. All data are expressed as mean ± SD, Welch’s *t*-test, n = 5. **(C)** SNX27^ΔIEC^ mice had greater disruption of crypt architecture and infiltration of inflammatory cells, overall contributing to a higher histological inflammation score. All data are expressed as mean ± SD, Welch’s *t*-test, n = 6. **(D)** Cytokine mRNA expression levels TNF-α and IL-6 were significantly higher in SNX27^ΔIEC^ mice compared to SNX27^Loxp^ mice. All data are expressed as mean ± SD, Welch’s *t*-test, n = 4-5. (**E)** AJs proteins Δ-catenin and E-cadherin were significantly downregulated in the colons of SNX27^ΔIEC^ mice compared to SNX27^Loxp^ mice. E-cadherin expression was also shifted from cell surface to cytoplasm with intestinal SNX27 deletion. Data are shown as mean ± SD, Welch’s *t*-test, n = 10. (**G)** Protein expression levels of Δ-catenin was analyzed via western blot showing significant degradation of Δ-catenin in SNX27^ΔIEC^ mice. Data are expressed as mean ± SD, Welch’s *t*-test, n = 4. All *p*-values are shown in the figures.

### SNX27 deletion disrupts cell-to-cell adhesion complex

Δ-catenin is a key player in maintaining cell-cell adhesion via its direct interactions with E-cadherin, thereby forming the adherens-junction (AJ) complex ^31^. This complex interaction is critical for the maintenance of the epithelial layer, as downregulation of AJ proteins has been shown to promote loss of epithelial cell polarity, epithelial-mesenchymal transition, and disruption of the epithelial barrier integrity ^32^. Therefore, we analyzed the protein expression levels of Δ-catenin and E-cadherin via immunofluorescence staining. We observed that SNX27^ΔIEC^ mice exhibited a significant loss of both Δ-catenin and E-cadherin from the membranes of the colonic epithelial cells **(Fig. 6E)**. We also observed a rearrangement in the localization of E-cadherin as it shifted from being expressed at the cell surface in SNX27^Loxp^ mice to being accumulated in the cytoplasm of SNX27^ΔIEC^ mice IECs. Furthermore, the western blot analysis showed a significant reduction of Δ-catenin protein level in the colon of SNX27^ΔIEC^ mice, compared to that in the SNX27^Loxp^ mice **(Fig. 6F)**.

### SNX27**^Δ^**^IEC^ mice are susceptible to DSS-induced colitis

We further hypothesized that SNX27^ΔIEC^ mice are more susceptible to colitis-associated intestinal damage ^25^. We challenged SNX27^Loxp^ and SNX27^ΔIEC^ mice with 5% DSS for 7 days to induce acute colitis **(Fig. 7A)**. We monitored the mice daily for bodyweight changes and observed that SNX27^ΔIEC^ mice significantly lost more bodyweight by the end of treatment **(Fig. 7B)**. Our histopathological analysis showed a higher infiltration of inflammatory cells and aberrant crypt architecture in the colon tissues of DSS treated SNX27^ΔIEC^ mice, which also correlated with a significant increase in histological inflammation score **(Fig. 7C)**. Accordingly, we observed that the DSS treated SNX27^ΔIEC^ mice had significant shortening of colon and small intestine, as compared to all other groups **(Fig. 7D and Fig. 7E)**. Using mice serum samples, we also analyzed intestinal permeability levels and observed a significantly higher concentration of FITC-Dextran in DSS treated SNX27^ΔIEC^ mice, compared to SNX27^Loxp^ mice treated with DSS **(Fig. 7F)**. Taken together, these data strongly suggest that lack of intestinal epithelial SNX27 promotes susceptibility towards DSS-induced colitis in mice resulting in aggravated intestinal tissue injury due to inflammation.

**Figure 7.**
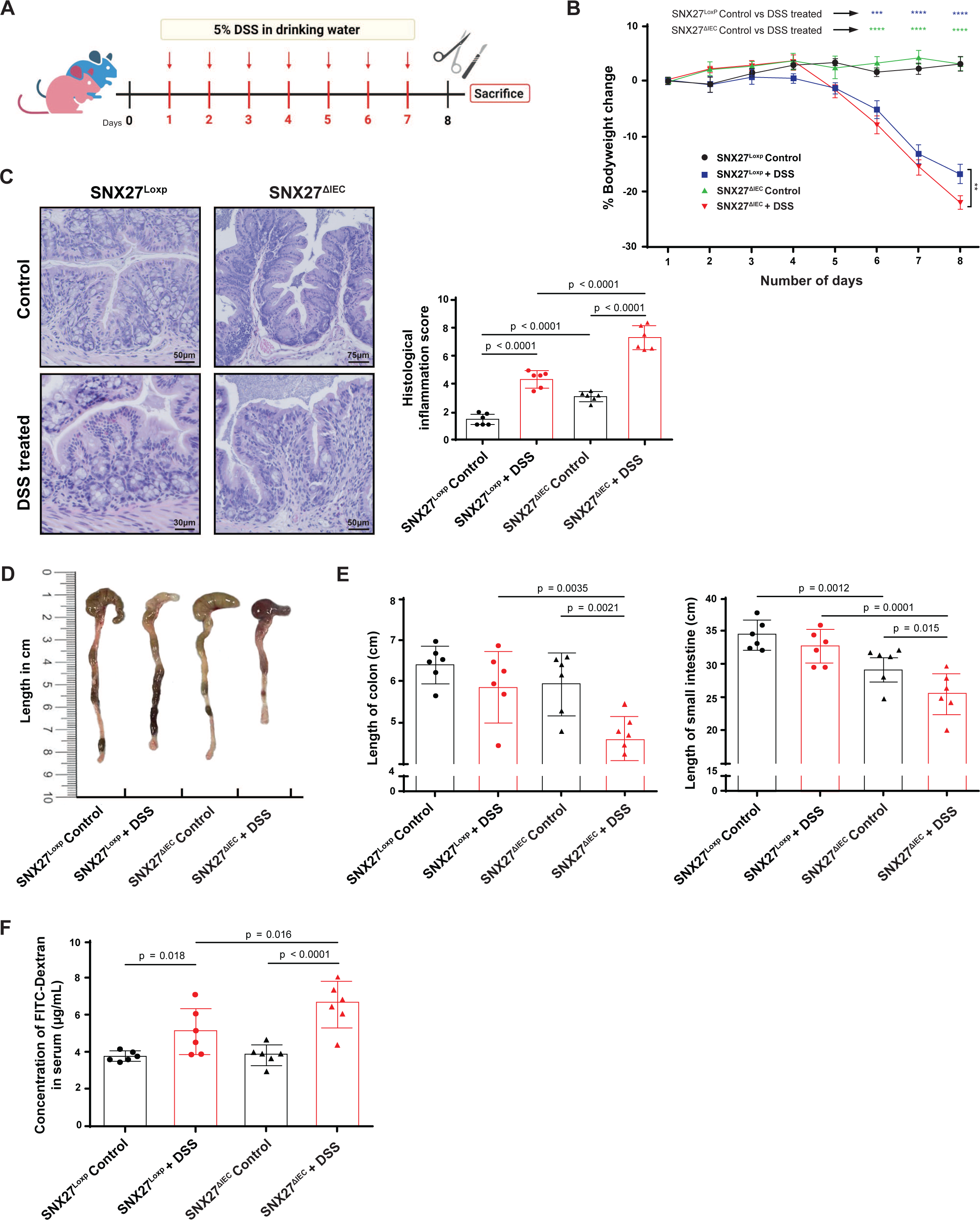
Intestinal epithelial SNX27 deletion enhanced susceptibility of mice towards intestinal tissue damage and inflammation during DSS-induced colitis. (A) Schematic overview of acute colitis mouse model. Mice were administered with 5% DSS in drinking water for 7 days (day 1-day 7) followed by sacrificing on day 8 for sample preparation. (B) Percentage body weight change revealed significantly greater loss of body weight in SNX27^ΔIEC^ mice challenged with DSS. All data are expressed as mean ± SD, two-way ANOVA test, n = 6. **(C)** H&E staining of colon tissues revealed a higher infiltration of inflammatory cells, aberrant crypt architecture and significant increase in histological inflammation score in the DSS treated SNX27^ΔIEC^ mice. All data are expressed as mean ± SD, one-way ANOVA test, n = 6. **(D)** and **(E)** SNX27^ΔIEC^ mice treated with DSS had significant shortening of colon and small intestine, as compared to all other groups. All data are expressed as mean ± SD, one-way ANOVA test, n = 6. **(F)** Intestinal permeability levels were higher in SNX27 ^ΔIEC^ mice treated with DSS compared to other groups, as determined by higher concentration of FITC-Dextran detected in serum. Data are expressed as mean ± SD, one-way ANOVA test, n = 6. All *p*-values are shown in the figures.

### Severity of DSS-induced colitis in SNX27**^Δ^**^IEC^ mice is correlated with increased disruption of cellular junction proteins at the epithelial barrier

As we observed loss of epithelial barrier proteins in SNX27^ΔIEC^ vs SNX27^Loxp^ mice, we further analyzed the extent of damage induced by DSS treatment on the expression and localization of TJ/AJ associated proteins in mice colons. Immunofluorescence (IF) revealed that ZO-1 expression was severely diminished in SNX27^ΔIEC^ mice treated with DSS among all the other control and treated groups **(Fig. 8A)**. Not only E-cadherin localization was re-arranged to the cytoplasm, but also its IF density was downregulated upon DSS treatment in SNX27^ΔIEC^ mice **(Fig. 8B)**. The protein expression level of Claudin10 was significantly lower between SNX27^ΔIEC^ and SNX27^Loxp^ mice, both at the basal level and upon treatment with DSS **(Fig. 8C)**.

**Figure 8.**
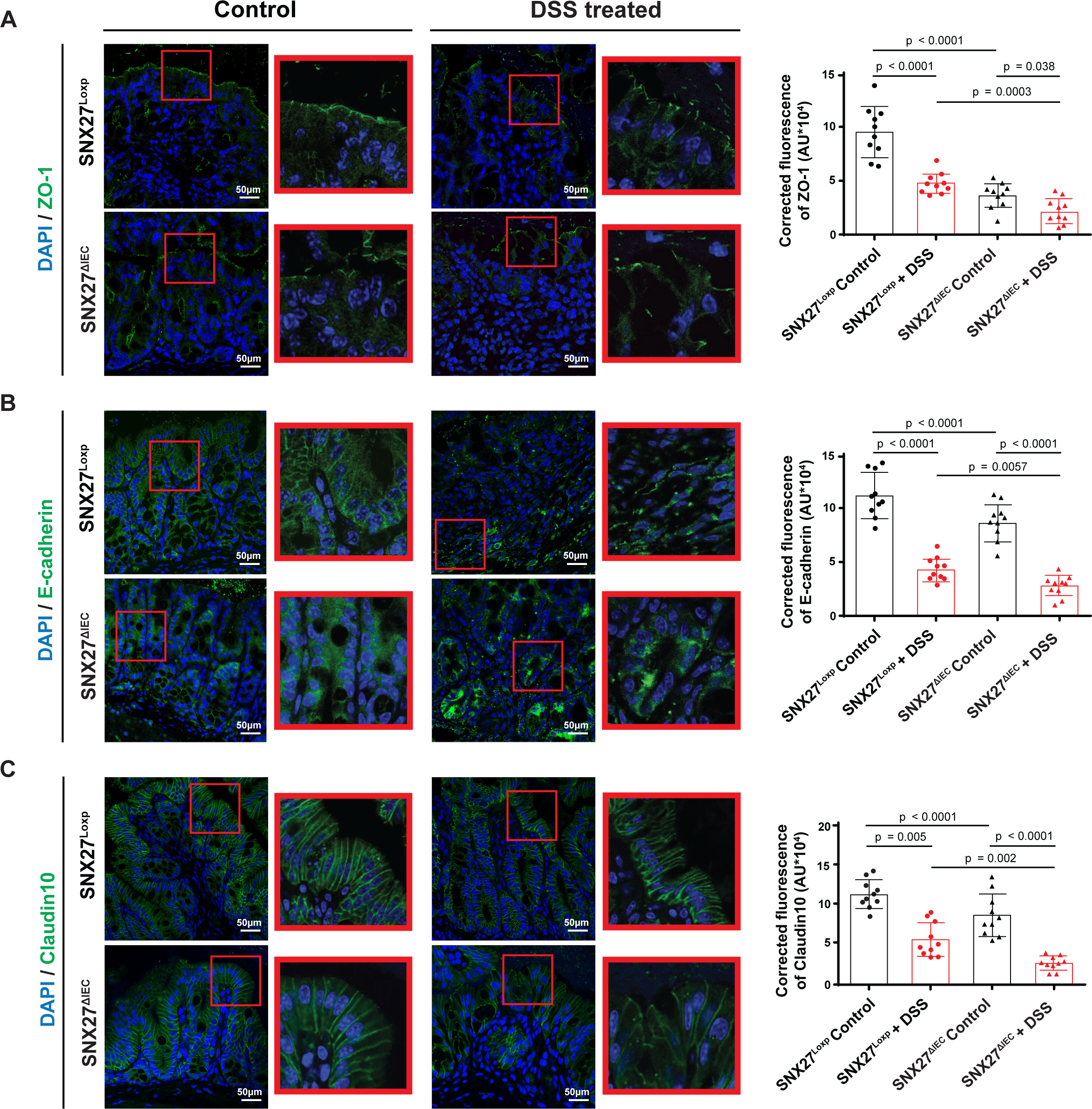
Severity of DSS-induced colitis in SNX27^ΔIEC^ mice was correlated with increased disruption of tight/adherens junction proteins at the epithelial barrier. **(A)** IF staining intensity and localization of ZO-1 at the epithelial barrier of proximal colon tissues was downregulated with DSS treatment, however the disruption was much greater in SNX27^ΔIEC^ mice at basal level as well as under colitis model, as compared to SNX27^Loxp^ mice. The corrected IF intensity was quantified with ImageJ. Data are expressed as mean ± SD, one-way ANOVA test, n = 10. **(B)** E-cadherin staining intensity in proximal colon tissues was diminished in the DSS treated groups, however the localization rearrangement from membrane to cytoplasm was more prominent in SNX27^ΔIEC^ mice. Data are expressed as mean ± SD, one-way ANOVA test, n = 10. **(C)** Immunofluorescence staining intensity of Claudin10 in proximal colon tissues is much lower in SNX27^ΔIEC^ mice at basal level as well as under colitis model, however the downregulation is further exacerbated with DSS treatment. Data are expressed as mean ± SD, one-way ANOVA test, n = 10. All *p*-values are shown in the figures.

**Figure 9.**
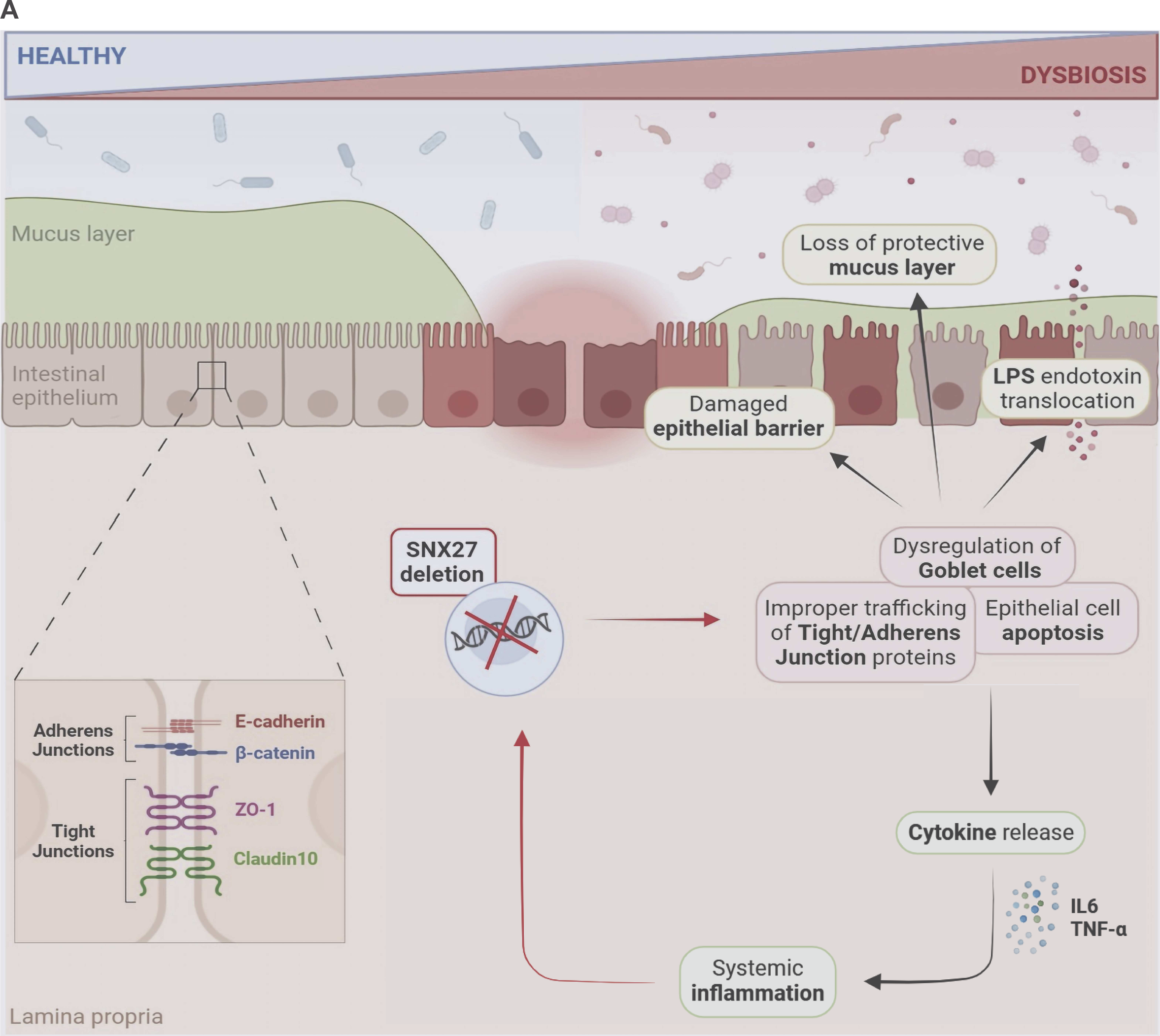
A working model of Impacts of intestinal epithelial SNX27 in chronic inflammation. Lack of intestinal epithelial SNX27 leads to gut leakiness and systemic inflammation due to dysfunction of epithelial junctions, potential dysbiosis, and proinflammatory cytokine release, thus increasing the susceptibility towards intestinal disorders, such as IBD.

## Discussion

In this study, we investigated the tissue specific roles of intestinal epithelial SNX27 in the context of IBD. Through the analysis of public datasets, we report significantly downregulated expression levels of SNX27 in patients with IBD, including those suffering from UC as well as CD. We found progressive loss of SNX27 in inflamed vs uninflamed tissues of IBD patients, suggesting its involvement in the intestinal inflammation. Using animal models, we further demonstrate that SNX27 plays a crucial role in regulating gut physiology. Deletion of SNX27 from IECs resulted in an overall imbalanced intestinal homeostasis, loss of mucus layer, and breakdown of epithelial barrier which highlights the mechanism through which SNX27^ΔIEC^ mice exhibit increased susceptibility towards colitis. Compared to Loxp mice, SNX27^ΔIEC^ mice showed significantly greater loss in bodyweight, more severe physiological distress symptoms, such as bloody diarrhea, and sensitivity towards DSS treatment. In summary, intestinal SNX27 conditional knockout exacerbated colitis severity due to poor barrier integrity and increased inflammatory damage. This is consistent with the observation of reduced SNX27 in IBD patients. Hence, these results highlight the importance of SNX27 in the intestinal tissue and shed light towards the previously unknown roles of SNX27 in the pathophysiology of human IBD.

SNX27 is critical for neurodevelopmental processes and whole-body knockout of SNX27 in mice has shown to be embryonically lethal ^20^. Our study is the first to report the physiological role of SNX27 in the intestines. Using our newly developed SNX27^ΔIEC^ mouse model, we found that loss of intestinal SNX27 induces drastic changes in development as SNX27^ΔIEC^ mice weighed significantly lower than SNX27^Loxp^ mice. We also observed significant attrition of small intestine and colon lengths in SNX27^ΔIEC^ mice, compared to the SNX27^Loxp^ mice. A previous *in vitro* study performed by Singh et al. has shown that the PDZ domain of SNX27 directly interacts with the C-terminus of sodium/proton exchanger-3 (NHE3), which regulates sodium reabsorption and hydrogen secretion. Interestingly, *in vitro* knockout of SNX27 disrupted the activity of NHE3, but did not change the total level of NHE3 protein ^33^. This suggests mechanisms though which deletion of intestinal SNX27 *in vivo* might induce nutrient malabsorption, which then leads to lighter body weights as seen in SNX27^ΔIEC^ mice. Interestingly, the weight of spleen was found to be heavier in SNX27^ΔIEC^ mice, which may indicate towards an active immune system. This is further reflected by a significantly higher inflammation score observed in the proximal colon tissues of SNX27^ΔIEC^ mice, compared to the SNX27^Loxp^ mice. It is crucial to note that even under normal growth conditions and lack of external stressors, we observed a greater presence of immune cells in the lamina propria, thickness of the submucosa, and distortion of crypt architecture upon deletion of intestinal epithelial SNX27, all of which represents SNX27’s tissue-specific protective role in mitigating inflammatory responses.

Interestingly, we observed a simultaneous increase in the rates of proliferation and apoptosis of IECs in the SNX27^ΔIEC^ mice. However, there were remarkable differences in the location of these proliferative and apoptotic cells. In a healthy state, the base of the intestinal crypts is a highly proliferative zone of progenitor cells while apoptosis occurs among the differentiated cells at the apical surface, thus constantly replenishing the intestinal epithelium. This observation is consistent in the SNX27^Loxp^ mice, however, we observed a significant increase in the number of apoptotic cells along the basolateral and migratory space of the crypts in SNX27^ΔIEC^ mice, compared to SNX27^Loxp^ mice. These findings are similar with those reported in patients with IBD, as studies have shown that IEC apoptosis increases in the crypt proliferative zones in both UC and CD, while also promoting mucosal ulcerations ^34^. Some studies have shown an increase in proliferative responses to counterbalance cell death and sustained epithelial injury in the colonic mucosa, such as upon DSS induced colitis in rats ^35^ and in patients with UC ^36^. We observed that SNX27^ΔIEC^ mice had significantly higher proliferation in the intestinal crypts as compared to SNX27^Loxp^ mice. This incidence of IEC hyperproliferation may be a defensive response to the excessive apoptosis to preserve intestinal homeostasis. However, abnormal proliferation rates are known to promote tumorigenesis, while persistent increase in cell death leads to breakdown of the epithelial barrier, facilitating mucosal invasion of pathogens and toxins, resulting in systemic inflammation and pathogenesis of IBD ^26^.

We observed a significant reduction in the numbers of Paneth and Goblet cells in SNX27^ΔIEC^ mice relative to SNX27^Loxp^ mice. We also observed poor differentiation in both types of secretory cells in the knockout mice. The imbalance between rates of IEC proliferation and apoptosis may alter progenitor cells differentiation and localization, however further studies are required to examine the mechanisms through which intestinal SNX27 may regulate different IEC lineages. Paneth cells secrete antimicrobial peptides that are essential for host defense against harmful pathogens and abnormality in the Paneth cells population has previously been reported in various intestinal disorders ^37^. Similarly, Goblet cells secrete mucins, forming a protective mucus barrier ^29^. This layer prevents unwanted gut luminal contents from coming into contact with the host intestinal environment, and loss of goblet cells has been shown to activate host immune responses and promote gut inflammation ^29^. Our data revealed that intestinal SNX27 deletion causes significant loss of the protective mucus layer, the first line of defense in the host gastrointestinal system. Therefore, deletion of SNX27 may increase the susceptibility towards intestinal tissue injury by promoting abnormal secretory cellular differentiation and loss of mucus layer, as observed in other colitis models ^38^.

We report in SNX27^ΔIEC^ mice a significant dysregulation of essential TJ proteins ZO-1 and Claudin10, as well as Δ-catenin and E-cadherin complex forming AJ. Right next to the mucus layer, the intestinal epithelial barrier forms the next protective wall defending the host system against invasion from harmful pathogens and toxins. Weakening of the epithelial barrier integrity increases intestinal leakiness and allows bacterial translocation into host tissues, further contributing to chronic inflammation and tissue injury ^25^. SNX27 is known to carry a PDZ domain and has already been reported to directly interact with ZO-2, which belongs to the same family of zonula occludens as ZO-1 ^39^. Here, we demonstrate a potential interaction that has not been reported before as evidenced by the correlation between loss of SNX27 and downregulation of ZO-1. Claudins are known to carry a PDZ-binding motif in their C-terminus which allows them to interact with ZO-1 and form the tight junctions ^19^. However, they also exhibit strong affinity towards other PDZ-domain carrying proteins, such as SNX27, which may explain our observations of highly dysregulated expression levels of Claudin10 resulting in weakening barrier integrity in SNX27^ΔIEC^ mice.

Consistent with previous literature, loss of SNX27 resulted in significant downregulation of Δ-catenin ^40^. Because the interaction between β-catenin and the cytoplasmic tail of E-cadherin stabilizes the AJ complex, loss of β-catenin from the plasma membrane also results in a downregulation of E-cadherin, which accumulates in the cytosol of IECs lacking SNX27. AJ proteins are crucial for maintaining the epithelial barrier as they facilitate strong cell-cell adhesion between adjacent IECs ^31, 32^. Therefore, intestinal SNX27 has the potential to mediate cell-cell adhesion by altering the protein expression levels and localization of critical AJ complex proteins at the intestinal epithelial barrier. β-catenin is a central player in the Wnt signaling pathway, where its phosphorylation status determines whether it is degraded or accumulates to activate downstream gene expression and proliferation ^41^. Previous studies have shown that proper regulation of β-catenin phosphorylation is crucial for maintaining a balance between tissue regeneration and inflammation in the intestine ^42^. Additionally, SNX27 is known to interact with Fzd7, a receptor and activator of the Wnt signaling pathway ^43^. What remains to be investigated, however, is the status of the excess free β-catenin in the intestinal epithelium. Therefore, further research is warranted to explore how SNX27-mediated regulation of β-catenin phosphorylation in IBD pathogenesis.

In the serum of SNX27^ΔIEC^ mice, we detected increased LPS, which is a key indicator of “leaky gut” and intestinal inflammation ^44^. LPS is a component of the outer membrane of Gram-negative bacteria which translocate from the intestinal lumen into the bloodstream when the epithelial barrier is compromised and intestinal permeability is increased ^44^. Serum cytokine levels showed significant upregulation of proinflammatory IL-6 and TNF-α release with reduced intestinal SNX27. Previous studies have reported that TNF-α and IL-6 have the potential to regulate various key inflammatory signaling pathways ^12^. This activation of systemic immune responses may facilitate further epithelial damage, feeding into the cycle of persistent barrier dysfunction and intestinal inflammation. Therefore, in our future studies we aim to investigate the effects of intestinal epithelial SNX27 deletion on the regulation of inflammatory signaling pathways commonly found to be aberrantly activated in human IBD ^45^.

In summary, basal level changes observed in SNX27^ΔIEC^ mice revealed a loss of intestinal epithelial barrier integrity due to dysregulated TJ/AJ proteins. Our study provides novel insights into the relationship between tissue-specific SNX27 expression and gut inflammatory mediators, such as Paneth/Goblet cell distribution, epithelial barrier proteins, etc. which highlight the complexity of SNX27-mediated intestinal physiology. Further investigations of intestinal SNX27 are warranted to understand its untapped potential as a therapeutic target to maintain intestinal integrity and inhibit chronic inflammation.

## Materials and Methods

### Human gene expression datasets

To determine the clinical relevance of SNX27 in human inflammatory bowel disease ^11^, including ulcerative colitis and Crohn’s disease ^12^, we analyzed publicly available microarray data uploaded in the NCBI Gene Expression Omnibus database (GEO) (https://www.ncbi.nlm.nih.gov/geo/). We determined the expression levels of SNX27 in IBD patient samples by identifying two different GEO database reference series. In the study with accession number GSE6731, comparisons were drawn between 13 UC, 19 CD and 4 healthy patients ^21^. Similarly, in another study with accession number GSE11223, colonic epithelial biopsies of various anatomic locations, including sites with and without active inflammation, from 67 patients with UC and 31 healthy controls were used to determine the differences in SNX27 expression ^22^.

### Animals

SNX27loxp/loxp (SNX27^Loxp^) mice were originally reported by Munoz et al ^46^. These mice, generously provided to us by Dr. Paul Slesinger, were developed by crossing heterozygous mice expressing knockout-first promoter line SNX27tm1a(KOMP)Wtsi (UC Davis KOMP Repository) to flp deleter expressing B6.SJL-Tg(ACTFLPe)9205Dym/J mice (JAX stock, 005703) ^46^. We further generated SNX27^ΔIEC^ mice carrying a conditional intestinal epithelial cell specific deletion of SNX27 by crossing SNX27^Loxp^ with Vil1/villin-Cre mice (Jackson Laboratory, 004586). All experiments were performed on 8-10 weeks old mice, including male and female. Animals were housed in the Biologic Resources Laboratory at the University of Illinois Chicago (UIC), provided with water ad libitum, and maintained in a 12 h dark/light cycle. All animals were utilized in accordance with the UIC Animal Care Committee and the Office of Animal Care and Institutional Biosafety guidelines under the approved animal protocol #ACC 21-177.

### Induction of colitis in mice

Mice at the age of 8-10 weeks old were randomly distributed into either control or treatment groups. Acute colitis was induced in the treatment group by supplementing 5% (w/v) dextran sodium sulphate (DSS) (MW = 40-50 kDa, MP Biomedicals) dissolved in drinking water provided ad libitum for 7 days along with a regular chow diet. All mice were monitored daily for bodyweight fluctuations, diarrhea, and/or bloody stool. At day 8, mice were humanely sacrificed under anesthesia and samples were harvested for downstream processing or storage.

### Immunoblotting

Mouse intestinal epithelial cells were harvested by gently scraping longitudinally dissected ileum and colon tissues, from proximal to distal end. The cells were collected in lysis buffer (10 mM Tris pH 7.4, 150 mM NaCl, 1 mM EDTA, 1 mM EGTA pH 8.0, 1% Triton X-100, 0.2 mM sodium ortho-vanadate, and protease inhibitor cocktail) and protein concentrations were measured using BioRad Reagent (BioRad), as described previously ^47^. Equal amounts of protein were separated by SDS-polyacrylamide gel electrophoresis, transferred to nitrocellulose membrane, blocked with 5% BSA, and incubated overnight at 4°C with desired primary antibodies as listed in Table 1. The following day, membranes were washed and incubated with target secondary antibodies for 1 hour and bands were visualized using Enhanced Chemiluminescence detection (ECL) reagent (Thermo Fisher Scientific).

**Table 1.**
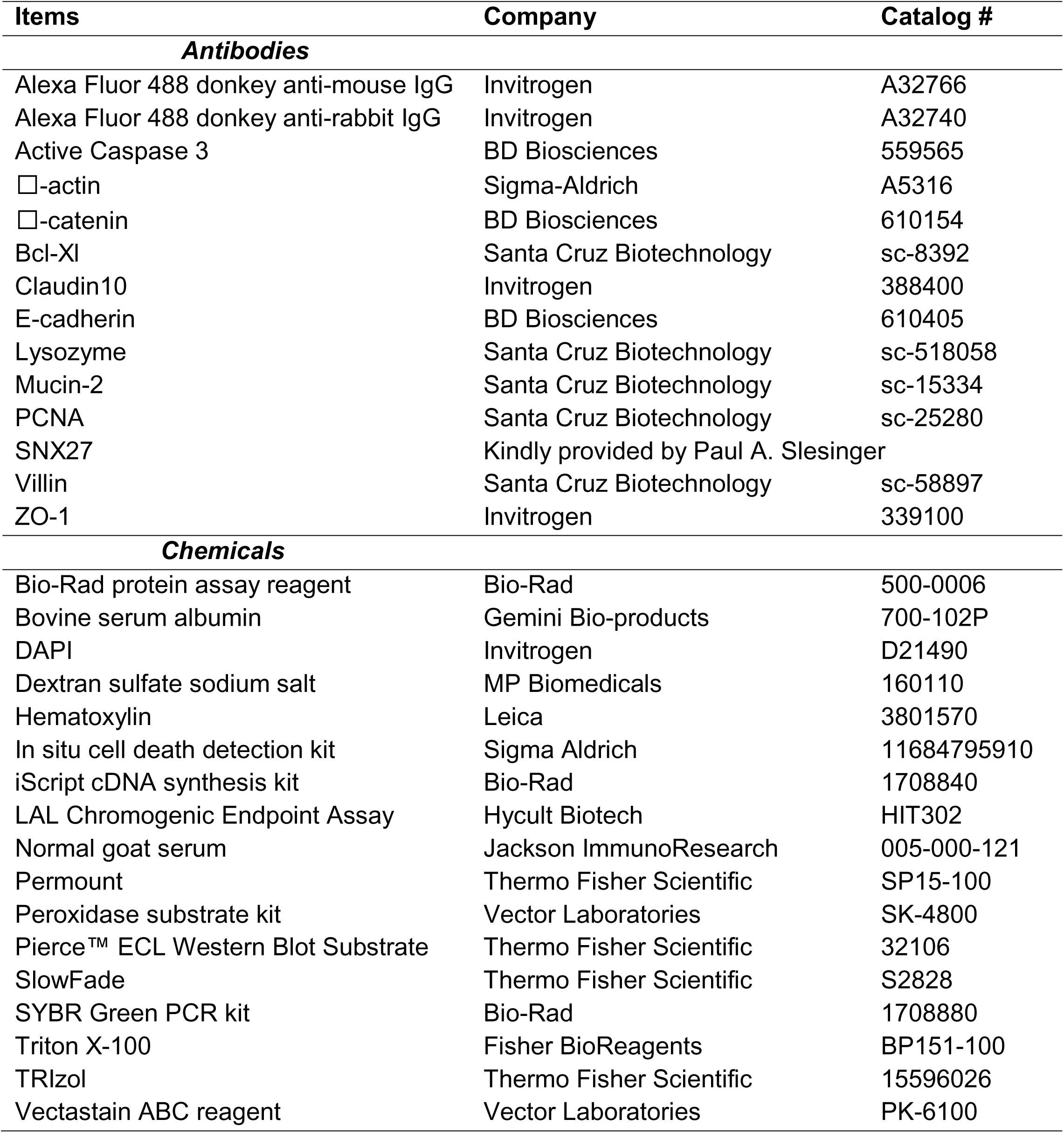
Methods key resources table.

### Histology of mice colon tissues

Mice ileum and proximal colon tissues were harvested, fixed in 10% formalin (pH 7.4), and paraffin-embedded. Slides of tissue sections were prepared at a thickness of 4 μm which were deparaffinized in xylene and rehydrated by passing through graded alcohol. Slides were then stained with hematoxylin and eosin followed by coverslip mounting with permount and were used to determine the histological damage and inflammation scores as previously described ^48^.

### Immunohistochemistry (IHC) staining

After preparation of the 4μm thick paraffin-embedded ileum and proximal colon sections on slides, antigen retrieval was achieved by incubating the slides for 15 min in hot preheated sodium citrate (pH 6.0) buffer followed by 30 min of cooling at room temperature. Endogenous peroxidases were quenched by incubating the slides in 3% hydrogen peroxide for 10 min, followed by three rinses with PBS. Slides were then incubated for 1 h with blocking buffer prepared with 2% bovine serum albumin (BSA) (Gemini Bio-products, 700-102P), 1% normal goat serum (Jackson ImmunoResearch Laboratories, 005-000-121), and 1% Triton X-100 (Sigma-Aldrich) in 1X-TBST to reduce nonspecific background. Primary antibodies as listed in Table 1 were applied overnight in a cold room. After three rinses with 1X-TBST, the slides were incubated with secondary antibody (1:100, Jackson ImmunoResearch Laboratories, Cat.No.115-065-174) for 1 h at room temperature. After washing with 1X-TBST for 10 min, the slides were incubated with vectastain ABC reagent (Vector Laboratories, Cat.No. PK-6100) for 1 h. After washing with 1X-TBST for five min, color development was achieved by applying a peroxidase substrate kit (Vector Laboratories, Cat.No. SK-4800) for 2 to 5 min, depending on the primary antibody. The duration of peroxidase substrate incubation was determined through pilot experiments and was then held constant for all of the slides. After washing in distilled water, the sections were counterstained with hematoxylin (Leica, Cat.No.3801570), dehydrated through ethanol and xylene, and cover-slipped using a permount (Fisher Scientific, Cat.No.SP15-100). The stained slides were analyzed using the ImageJ Fiji software for Semi-quantification, as described before ^49^.

### Immunofluorescence (IF) staining

Mouse ileum and proximal colon tissues were freshly harvested and fixed in 10% neutral buffered formalin followed by paraffin-embedding. Tissue sections were then cut at a thickness of 4μm and prepared on slides for immunofluorescence labelling, as described previously ^50^.

Slides were incubated for 1 h with blocking buffer prepared with 2% bovine serum albumin (BSA) (Gemini Bio-products, 700-102P), 1% normal goat serum (Jackson ImmunoResearch Laboratories, 005-000-121), and 1% Triton X-100 (Sigma-Aldrich) in 1X-TBST to reduce nonspecific background. After overnight incubation at 4°C with desired primary antibodies, as listed in Table 1, tissue sections were then labelled with the secondary anti-mouse or anti-rabbit Alexa Fluor 488 and DAPI for 1 hour at room temperature. Lastly, the slides were coverslip mounted with SlowFade (Thermo Fisher Scientific, S2828) and the edges were sealed to prevent from drying. The fluorescence intensity was visualized under a Zeiss laser scanning microscope LSM 710 (Carl Zeiss Inc.) followed by quantitative analysis using ImageJ software.

### Quantification of Paneth cells in the small intestine

Paneth cell staining was performed via immunofluorescence using the anti-lysozyme antibody. The morphological changes and varying expression of lysozyme were characterized as - D0 (normal PCs) and D1 to D3 (abnormal PCs), as described in previous publications ^28^.

### Terminal deoxynucleotidyl transferase dUTP nick end labeling (TUNEL) staining

The number of apoptotic cells were determined using the In Situ Cell Death Detection Kit (Sigma-Aldrich, 11684795910) in 10% neutral formalin-fixed paraffin-embedded 4μm thick proximal colon tissue sections. Briefly, antigen retrieval was achieved after deparaffination and rehydration by incubating the slides for 15 min in hot preheated sodium citrate (pH 6.0) buffer. Then the slides were cooled down to room temperature and washed with PBS for 10 minutes. After blocking, the slides were incubated with a TUNEL Reaction Mixture for 1 hour at 37°C. The tissues were then mounted with SlowFade Antifade Kit (Thermo Fisher Scientific, S2828) and the staining was examined with a Zeiss laser scanning microscope LSM 710 (Carl Zeiss Inc.).

### Fluorescence in-situ hybridization (FISH) staining

Fluorescence in situ hybridization (FISH) ^30^ was performed using antisense single-stranded DNA probes targeting the bacterial 16S ribosomal RNA. The EUB338 (5′-GCTGCCTCCCGTAGGAGT-3′) conjugated to Alexa Fluor488 probe was used for bacteria as 16S ribosomal RNA is highly conserved between different species of bacteria. FISH was performed on 4 μm thick paraffin-embedded sections of ileal and colonic mucosal biopsies fixed in Carnoy’s fixative solution. Tissue sections were deparaffinized in xylene, dehydrated with 100% ethanol, dried and incubated in 0.2 M HCl for 20 min. Then, the slides were heated in 1 M NaSCN at 80°C for 10 min after being washed with saline sodium citrate (SSC) solution. Tissues were pepsin-digested (4% pepsin in 0.01 N HCl) for 20 min at 37°C and incubated with 10% buffered formalin for 15 min. The tissues were washed and dried and hybridized with the probes (5 ng/μL) in hybridization buffer (0.9 mol/L NaCl, 30% formamide, 20 mmol/L Tris–HCl, and 0.01% SDS in SSC solution) at 96°C for 5 min and then at 37°C overnight. For visualization of the epithelial cell nuclei, the slides were counterstained with DAPI/ antifade solution. Slides were examined with a Zeiss laser scanning microscope LSM 710 (Carl Zeiss Inc.) and fluorescence staining intensity was determined using ImageJ software.

### Intestinal permeability assay

Mice were supplemented with 50mg/g of Fluorescein Dextran (MW = 3 kDa, Sigma-Aldrich) diluted in HBSS via oral gavage. Four hours later, blood samples were collected to measure fluorescence intensity, as previously reported ^51^.

### Serum LPS detection

LPS in serum samples was measured with limulus amebocyte lysate (LAL) chromogenic end point assay (Hycult Biotech, HIT302) according to the manufacturer’s indications. The samples were diluted 1:4 with endotoxin-free water and then heated at 75°C for 5 min on a warm plate to denature the protein before the reaction. A standard curve was generated and used to calculate the concentrations, which were expressed as EU/mL, in the serum samples.

### Real-time PCR analysis

Total RNA was extracted from mice proximal colon tissues using TRIzol reagent (Thermo Fisher Scientific), according to manufacturer’s protocol. RNA reverse transcription was performed using the iScript complementary DNA synthesis kit (Bio-Rad), as per manufacturer’s instructions. Then, the cDNA products were subjected to quantitative real-time PCR using the iTaq Universal SYBR green supermix (Bio-Rad) and primers from Primer Bank (Cambridge) as listed in Table 2. All mRNA expression levels were normalized to Δ-actin levels of the same sample. The percentage expression was calculated as the ratio of the normalized value of each sample to that of the corresponding untreated control samples. All real-time PCR reactions were performed in triplicates.

**Table 2.**
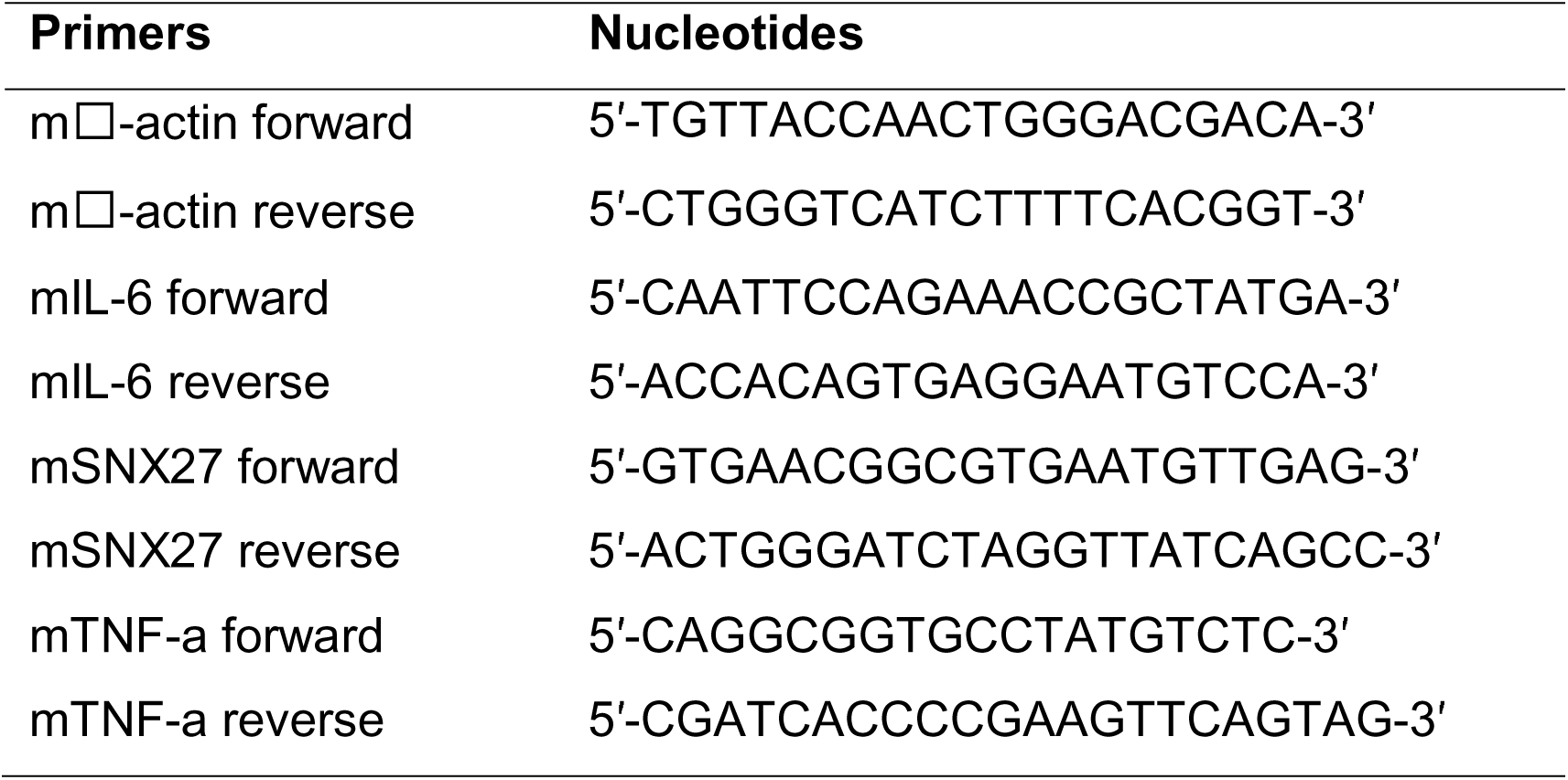
Real-time PCR primers.

### Statistical analysis

All experiments were repeated at least three times and data represented in bar graphs are expressed as mean ± SD or ± SEM. All performed statistical tests were 2-sided and p-values of 0.05 or less were considered statistically significant. Comparisons between two groups with normal distribution were analyzed by Welch’s *t*-test. Comparisons between three or more groups were analyzed by either one-way or two-way ANOVA, as appropriate. To correct for multiple comparisons in ANOVA, *p*-values were adjusted by Tukey’s or Sidak’s method ^52^. All statistical analyses were performed using GraphPad Prism version 8.0.0 for Windows.

## Acknowledgements/Funds

We would like to acknowledge the UIC Cancer Center, the NIDDK/National Institutes of Health grant R01 DK134343, R01DK114126, Crohn’s & Colitis Foundation Senior Research Award (Grant No. 902766), and VA Merit Award VA 1 I01 BX004824-01 to Jun Sun. The study sponsors play no role in the study design, data collection, analysis, and interpretation of data. The contents do not represent the views of the United States Department of Veterans Affairs or the United States Government. We thank Dr. Paul A. Slesinger for sharing SNX27^Loxp^ mice and Duncan Claypool (MD/PHD student at UIC) for proofreading the manuscript.

## Authors’ contributions

SD: acquisition, analysis, and interpretation of data; drafting the manuscript; and statistical analysis. YZ: assistance with animal models and data analysis. YX: statistical analysis and manuscript drafting. JS: study concept and design, writing the manuscript, and obtained funding.

## Abbreviation list

AJ: Adherens junction
ANOVA: Analysis of variance
Bcl-xL: B cell lymphoma-extra large
BSA: Bovine serum albumin
CD: Crohn’s disease
DAB: Diaminobenzidine
DAPI: Diamidino-2-phenylindole
DNA: Deoxyribonucleic acid
DSS: Dextran sulfate sodium
ECL: Enhanced chemiluminescence
FERM: 4.1/ezrin/radixin/moesin
FISH: Fluorescence in situ hybridization
FITC: Fluorescein isothiocyanate
GEO: Gene expression omnibus
GC: Goblet cell
GFP: Green fluorescent protein
HBSS: Hanks balanced salt solution
H&E: Hematoxylin & Eosin
IBD: Inflammatory bowel disease
IEC: Intestinal epithelial cells
IHC: Immunohistochemistry
IF: Immunofluorescence
IL-6: Interleukin 6
LPS: Lipopolysaccharides
mRNA: Messenger ribonucleic acid
Muc2: Mucin-2
MW: Molecular weight
PBS: Phosphate buffered saline
PC: Paneth cell
PCNA: Proliferating cell nuclear antigen
PX: Phox homology
PDZ: Post-synaptic density 95/discs large/zonula occludens-1
qPCR: Quantitative polymerase chain reaction
RNA: Ribonucleic acid
siRNA: Small interfering ribonucleic acid
SNX: Sorting nexin
SSC: Saline Sodium Citrate
TBST: Tris-buffered saline with 0.1% Tween 20
TJ: Tight junctions
TNF-α: Tumor necrosis factor α
TUNEL: Terminal deoxynucleotidyl transferase dUTP nick end labeling
UIC: University of Illinois Chicago
UC: Ulcerative colitis
ZO-1: Zonula ocludens-1

